# Using genetic drug-target networks to develop new drug hypotheses for major depressive disorder

**DOI:** 10.1101/304113

**Authors:** Héléna A Gaspar, Zachary Gerring, Christopher Hübel, Christel M Middeldorp, Eske M Derks, Gerome Breen

**Author notes:** postal address: Social, Genetic and Developmental Psychiatry Centre, Institute of Psychiatry, Psychology and Neuroscience – PO80, De Crespigny Park, Denmark Hill, London, United Kingdom, SE5 8AF.

## Abstract

The major depressive disorder (MDD) working group of the Psychiatric Genomics Consortium (PGC) has published a genome-wide association study (GWAS) for MDD in 130,664 cases, identifying 44 risk variants. We used these results to investigate potential drug targets and repurposing opportunities. We built easily interpretable bipartite drug-target networks integrating interactions between drugs and their targets, genome-wide association statistics and genetically predicted expression levels in different tissues, using our online tool Drug Targetor (drugtargetor.com). We also investigated drug-target relationships and drug effects on gene expression that could be impacting MDD. MAGMA was used to perform pathway analyses and S-PrediXcan to investigate the directionality of tissue-specific expression levels in patients vs. controls. Outside the major histocompatibility complex (MHC) region, 25 druggable genes were significantly associated with MDD after multiple testing correction, and 19 were suggestively significant. Several drug classes were significantly enriched, including monoamine reuptake inhibitors, sex hormones, antipsychotics and antihistamines, indicating an effect on MDD and potential repurposing opportunities. These findings require validation in model systems and clinical examination, but also show that GWAS may become a rich source of new therapeutic hypotheses for MDD and other psychiatric disorders that need new – and better – treatment options.

## Introduction

There is an urgent need for new drugs to better treat major depressive disorder (MDD), with new modes of action as well as fewer side effects. The Psychiatric Genomics Consortium (PGC) has conducted a genome-wide association study (GWAS) of more than 130,664 MDD and broader depression cases and 330,470 controls identifying 44 loci associated with depression.^1^ Much new biology is suggested by these findings and we hypothesize that the collection of loci discovered by GWAS may have the potential to restart largely paused drug development pipelines. This is not without considerable technical challenges. At the moment, time-consuming manual assessment by expert biologists and geneticists is required for each GWAS locus. Analyzing all genome-wide results together may allow better prioritization of potential drug or therapeutic hypotheses^2,3^.

GWAS associations between single nucleotide polymorphisms (SNPs) and MDD can be used to assess the association of each gene or sets of genes, such as those defined by biological pathways. Pathway analysis has also been used to suggest new drug hypotheses by mapping drugs to the proteins they bind, and defining the sets of genes that encode the proteins as “drug gene-sets” whose association with a phenotype of interest can be estimated.^2, 4^ This process is a type of drug repositioning analysis aimed at finding potential new uses for existing drugs.^2^ In this paper, we propose to mine drug-protein/gene interactions from two main sources: drug-target relationships or activity profiles and drug effects on gene expression or perturbagen signatures”.^5^ Activity profiles can be derived from several databases such as PubChem BioAssays^6^ or ChEMBL,^7^ while the main source for perturbagen signatures is the CMAP database.^5^ Instead of using these resources separately, they can be used together to identify relevant drugs. However, simply generating the association between drug gene-sets and phenotypes is not enough; each gene-set is a subnetwork with different interaction types between drugs and proteins. Visualising these interactions could allow better and more rapid prioritization of drug gene-sets.

For this purpose, it may be useful to translate both activity profiles and perturbagen signatures into bipartite drug-target interaction networks. These can be constructed by linking drug nodes to targets nodes where the links or edges represent the type of drug-target interaction. Maggiora et al.^8^ suggested that these networks could be used to assess drug polypharmacology – the ability of drugs to interact with several targets – as well as target polyspecificity – the ability of targets to exhibit affinity towards multiple dissimilar molecular compounds.

In this paper, we build drug-target networks relevant to a given phenotype (MDD), by using the results from a well-powered PGC MDD GWAS for imputation of tissue-specific expression levels in patients vs. controls, and to generate genetic associations of known drug targets with MDD. We also present a network visualization tool – available at drugtargetor. com – which provides the opportunity to build networks linking these genetic data with a large number of drugs and drug classes, allowing detailed assessment of drug action possibly impacting MDD.

## Materials and Methods

### Genome-wide association study of major depressive disorder

The PGC MDD phase 2 analysis^1^ was a combined analysis of an anchor cohort of traditionally ascertained MDD cases (16,823 MDD cases and 25,632 controls)^1^ and an expanded cohort of more diversely assessed depression cases (113,841 MDD cases and 304,838 controls). A combination of polygenic scoring and linkage disequilibrium (LD) score genetic correlation comparisons between the anchor and expanded cohorts and samples showed strong evidence for genetic homogeneity between these groups.^1^ SNPs, insertions and deletions were imputed using the 1000 Genomes Project multi-ancestry reference panel.^10^ Association analyses were performed within each cohort using imputed marker dosages and principal components as covariates to account for population stratification. Principal components analysis was used to determine ancestry from genotyped SNPs.^11^ Summary statistics for 10,468,942 autosomal SNPs were then available for the analyses we present.

### Gene-based test of association

We used MAGMA v1.06^12^ to perform a gene-based test of association with the MDD GWAS summary statistics. Briefly, MAGMA generates gene-based p-values by combining adjacent SNP-based p-values within a defined gene window while accounting for LD. SNPs were mapped to genes if they were located 35 kb upstream or 10 kb downstream of a gene body including regulatory regions, and the gene p-value is obtained using the “multi=snp-wise” option, which aggregates mean and top SNP association models. A Bonferroni p-value threshold of 2.63 × 10^−6^, accounting for 19,079 ENSEMBL genes, was used to account for multiple testing. We used 1000 Genomes European data phase 3 as the reference LD set^10^.

### Transcriptome-wide association

To assess the impact of genetic variation underlying MDD on gene expression, we performed a transcriptome-wide association study (TWAS) using the S-PrediXcan software^13^. This approach estimates gene expression weights by training a linear prediction model in samples with both gene expression and SNP genotype data. The weights are then used to predict gene expression from GWAS summary statistics, while incorporating the variance and covariance of SNPs from a LD reference panel. We used pre-computed gene expression weights for 10 brain tissues (anterior cingulate cortex, caudate nucleus, cerebellar hemisphere, cerebellum, cortex, frontal cortex, hippocampus, hypothalamus, nucleus accumbens, and putamen) generated from the Genotype-Tissue Expression (GTEx) Consortium,^14^ and whole blood using the Depression Genes and Networks (DGN) cohort.^15^ The 1000 Genomes European data phase 3 was used as the reference LD set.^10^ These data were processed with beta values and standard errors from the MDD GWAS summary statistics to estimate the expression-GWAS association statistic. A transcriptome-wide significance threshold of *P* = 1.25 × 10^−6^, adjusting for all GTEx brain tissue and DGN associations (Bonferroni correction 0.05/39,936), was used to adjust for multiple testing.

### Definition of the druggable genome

We used 4,479 genes estimated to be “druggable” by Finan et al.^16^ (henceforth referred to as the “druggable genome”); they divided these genes into three “tiers” based on their importance in pharmaceutical development: tier 1 (targets of approved/clinical trial drugs), tier 2 (similar to tier 1 proteins or targeted by drug-like molecules), and tier 3 (proteins with lower similarity to tier 1 proteins, secreted or extracellular proteins, main druggable families). In the gene-based tests of association, genes were investigated regardless of their druggable status; however, we only used druggable genes to build drug-target networks. Information on genes with human or mouse phenotypes were also collected from the human-mouse disease connection database (HDMC), which gathers mouse data from Mouse Genome Informatics database (MGI)^17^ and human data from the National Center for Biotechnology Information (NCBI) and Online Mendelian Inheritance in Man (OMIM).^18^

### Definition of drug-target and drug-gene interactions

We collected two types of drug interactions: activity profiles (drug-target interactions) and perturbagen signatures (drug-gene interactions). Drug-target interactions are defined as any type of interaction between a drug and a protein target. Drug-gene interactions are changes in gene expression induced by a drug. We built an *annotation* dataset using interaction profiles from the drug-gene interaction database DGIdb v2.0,^19^ ChEMBL v.23^7, 20^, the psychoactive drug-gene database PDSP K_i_ DB, PHAROS,^21^ NCBI PubChem BioAssay,^22^ and DSigDB^16, 23^ (downloaded in June 2017), which also contains CMAP data. We subset experimental data from the *annotation* dataset to generate a more reliable *curated dataset*, discarding data from textmining approaches. The broad *annotation* set was used for all analyses and to annotate drug-target networks, while the *curated* subset was used to check which drug classes were enriched when restricting analyses to experimental data. A description of the bioactivities curation approach is provided in **Supplementary Text 1**.

### Enrichment of drug gene-sets and therapeutic classes

Approved drugs and their Anatomical Therapeutic Chemical (ATC) codes were identified by mapping all drug names to their PubChem compound identifier (CID) using the PubChem synonym database (ftp.ncbi.nlm.nih.gov/pubchem/Compound/Extras/CID-Synonym-filtered.gz), then mapping each CID to the corresponding ATC codes. The drugs were merged by ATC name, which could correspond to several CID entries and ATC codes. Each drug was then mapped to a gene-set using the collected drug-gene and drug-target interactions, and assigned a p-value generated by competitive pathway analysis (MAGMA), assessing the association between drug gene-set and phenotype. For the **annotation** set, 1946 drugs corresponding to 1738 individual gene-sets were tested; for the more reliable *curated* set, 1547 drugs were mapped to 1282 gene-sets. For each ATC hierarchical level, enrichment curves were drawn by ranking drug gene-sets by their association with MDD. The area under the enrichment curve (AUC) and associated p-value from Wilcoxon rank tests were used to evaluate the enrichment of drug classes for both *annotation* and *curated* sets separately.

### Bipartite drug-target networks

Bipartite drug-target networks were built using our online tool Drug Targetor (drugtargetor.com), presented here for the first time. The tool builds networks using two input files: a drug table with drug-phenotype associations and a target table with target-phenotype associations (cf. **Figure 1**). The drug-phenotype associations were obtained using the MAGMA and S-PrediXcan results, and the target-phenotype interactions were collected as described in the data collection section (cf. “Definition of drug-target and drug-gene interactions”). The networks are comprised of drug nodes and target nodes, the edges of which are connected based on the type of interaction. Drug Targetor defines nine types of drug-target interactions: increasing gene expression, decreasing gene expression, mixed (increasing or decreasing) gene expression, agonist/activator/positive allosteric modulator, partial agonist, antagonist/inhibitor/negative allosteric modulator, modulator (neither negative nor positive), inverse agonist, and mixed bioactivities (unknown or both agonist and antagonist). The drug nodes and target nodes are respectively ordered by decreasing association with MDD in -log_10_(P) units, where P is the pathway analysis p-value for drugs (MAGMA pathway analysis) and the gene-level p-value for targets (MAGMA gene-wise analysis). The predicted effects on gene expression in ten brain regions and whole blood are provided for each target.

**Figure 1.**
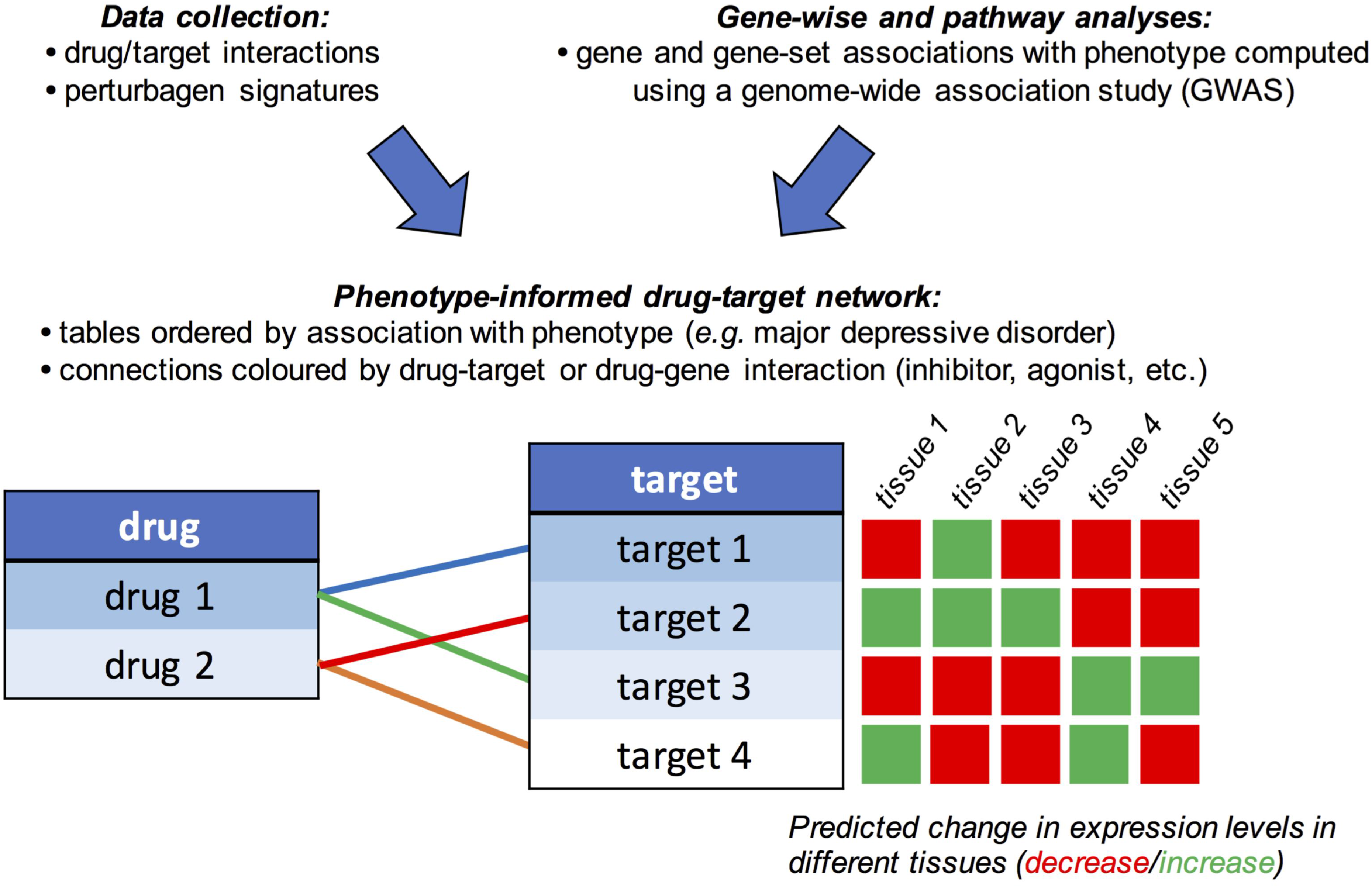
Workflow to build phenotype-informed bipartite drug-target networks, illustrating the principle behind the online tool “Drug Targetor” (drugtargetor.com).

## RESULTS

### Gene-based tests of association

We used MAGMA to map SNP-level association to individual genes, and filtered the data based on druggability (cf. “Definition of the druggable genome”). A total of 41 druggable genes achieved genome-wide significance (MAGMA *P* < 2.63 × 10^−6^), of which 25 were located outside the major histocompatibility complex (MHC) region (Table 1), and a further 21 genes (19 outside the MHC region) had suggestive significance (*P* < 2.63 × 10^−5^) (cf. **Supplementary Tables 1-2**).

**Table 1.** Druggable genes outside the major histocompatibility complex significant or suggestive in major depressive disorder. The -log_10_(p) column indicates the significance level as computed by MAGMA, the DGN whole blood and GTEx brain regions columns indicate the predicted change in expression level in the corresponding tissue.

| Gene name | $-\log_{10}(p)$ | DGN whole blood | GTEx brain regions |
| --- | --- | --- | --- |
| *NEGR1* | 16.63 | + (***) | +(1)/-(1) |
| *OLFM4* | 15.06 | + | +(2) |
| *CACNA1E* | 10.90 | - |  |
| *ESR2* | 9.32 | - (***) | -(1) |
| *PXDNL* | 8.88 | - |  |
| *DRD2* | 8.86 |  | +(1)/-(1) |
| *CHADL* | 8.36 | + | -(1) |
| *VRK2* | 8.32 | + | +(2) |
| *EP300* | 8.07 | + | -(1) |
| *CACNA2D1* | 8.00 |  |  |
| *RPS6KL1* | 7.81 | + | +(1)/-(1) |
| *GRM5* | 7.50 |  | +(2)/-(2) |
| *EMILIN3* | 7.41 |  | -(2) |
| *ENOX1* | 7.35 |  | -(1) |
| *HP* | 6.97 | - | +(6) |
| *PCDHA2* | 6.88 |  | +(1)/-(3) |
| *KCNB1* | 6.79 |  | -(1) |
| *GRIK5* | 6.41 | - | +(2)/-(2) |
| *FEN1* | 6.40 | - | -(1) |
| *PCSK5* | 6.15 | + | -(1) |
| *LINGO1* | 6.08 |  | +(5)/-(2) |
| *HSPD1* | 5.98 | - |  |
| *WDR1* | 5.75 | + |  |
| *TOP1* | 5.75 |  |  |
| *KMO* | 5.62 | - |  |
| *HTR1D* | 5.44 |  | +(4) |
| *MARK3* | 5.38 | - | +(3) |
| *DHODH* | 5.24 | - | +(2)/-(4) |
| *NCOR2* | 5.15 | + | +(1)/-(1) |
| *KLHL32* | 5.01 | - | +(1)/-(1) |
| *HDAC10* | 4.97 |  | +(2) |
| *CACNA1H* | 4.93 | + | +(3) |
| *COL8A2* | 4.92 | - | +(1)/-(2) |
| *MARK2* | 4.84 | - | +(1)/-(1) |
| *PPIF* | 4.81 | - | -(1) |
| *ADK* | 4.81 | + | -(1) |
| *MAPK12* | 4.78 | - | +(8)/-(1) |
| *CRB1* | 4.77 |  | -(1) |
| *RSPO1* | 4.73 |  | +(1) |
| *ABCC2* | 4.71 | - | -(1) |
| *GRM8* | 4.68 |  |  |
| *MAPK11* | 4.66 | - | +(1)/-(1) |
| *NTRK2* | 4.65 |  |  |
| *OTOA* | 4.62 |  | +(1) |

*** Bonferroni-significant S-PrediXcan results; +(1): predicted upregulation in one brain region; -(1): predicted downregulation in one brain region.

To gain insight into the potential functional consequences of DNA sequence variation underlying MDD, we imputed gene expression using S-PrediXcan. Overall, 35 protein-coding genes (16 outside the MHC) were significantly up-or downregulated in whole blood or brain (**Supplementary Table 3**). In whole blood, we found a significant association between MDD and the expression of two druggable genes outside the MHC region: *NEGR1* (*Z* = 7.35, *P* = 2.03 × 10^−13^) and *ESR2* (*Z* = -5.43, *P* = 5.66 × 10^−8^). The expression of two additional MHC druggable genes in whole blood -*BTN1A1* (*Z* = -5.91, *P* = 3.38 × 10^−9^) and *BTN3A2* (*Z* = 5.31, *P* = 1.12 × 10^−7^)- was also associated with MDD. Within GTEx brain tissues, the expression of the MHC druggable genes *BTN3A2* (upregulation in all brain regions), its paralog BTN3A3 (downregulation in cerebellar hemisphere), and *HIST1H4I* (downregulation in anterior cingulate cortex and nucleus accumbens) were significantly associated with MDD. We also found nominal evidence (cf. **Supplementary Tables 1-2**) for the upregulation of *DRD2* and *NEGR1* in brain cortex, *HTR1D* in the caudate and cortex, *MARK3* in the cerebellar hemisphere and hippocampus, downregulation of LINGO1 in the cerebellum, and *DHODH* in the frontal cortex. S-PrediXcan predictions in different tissues are not always concordant – for example, MDD is associated with decreased MARK3 expression in whole blood and increased expression in brain regions.

### Drug classes and their drug-target networks

We tested for the enrichment of MDD GWAS association signals within major therapeutic classes defined by ATC code. To correct for multiple testing and allow us to explore more hypotheses, we used the Benjamini and Hochberg false discovery rate (FDR)^24^ to adjust p-values. A total of 19 drug classes (FDR q-value < 0.05; **Figure 2** and **Supplementary Table 4**) were enriched for MDD GWAS association signals, of which five remained FDR-significant after restricting the analyses to experimentally validated drug-target and drug-gene interaction data (*curated set*). These included targets of psycholeptics (ATC code N05, *P* = 4.05 × 10^−7^), antipsychotics (N05A, *P* = 7.44 × 10^−6^), and non-selective monoamine reuptake inhibitors (N06AA, *P* = 6.72 × 10^−4^), highlighting the potential utility of MDD GWAS data for drug compound discovery and repositioning, but also targets of antihistamines (R06A, *P* = 4.67 × 10^−6^) and sex hormones and modulators of the genital system (G03, *P* = 1.98 × 10^−3^).

**Figure 2.**
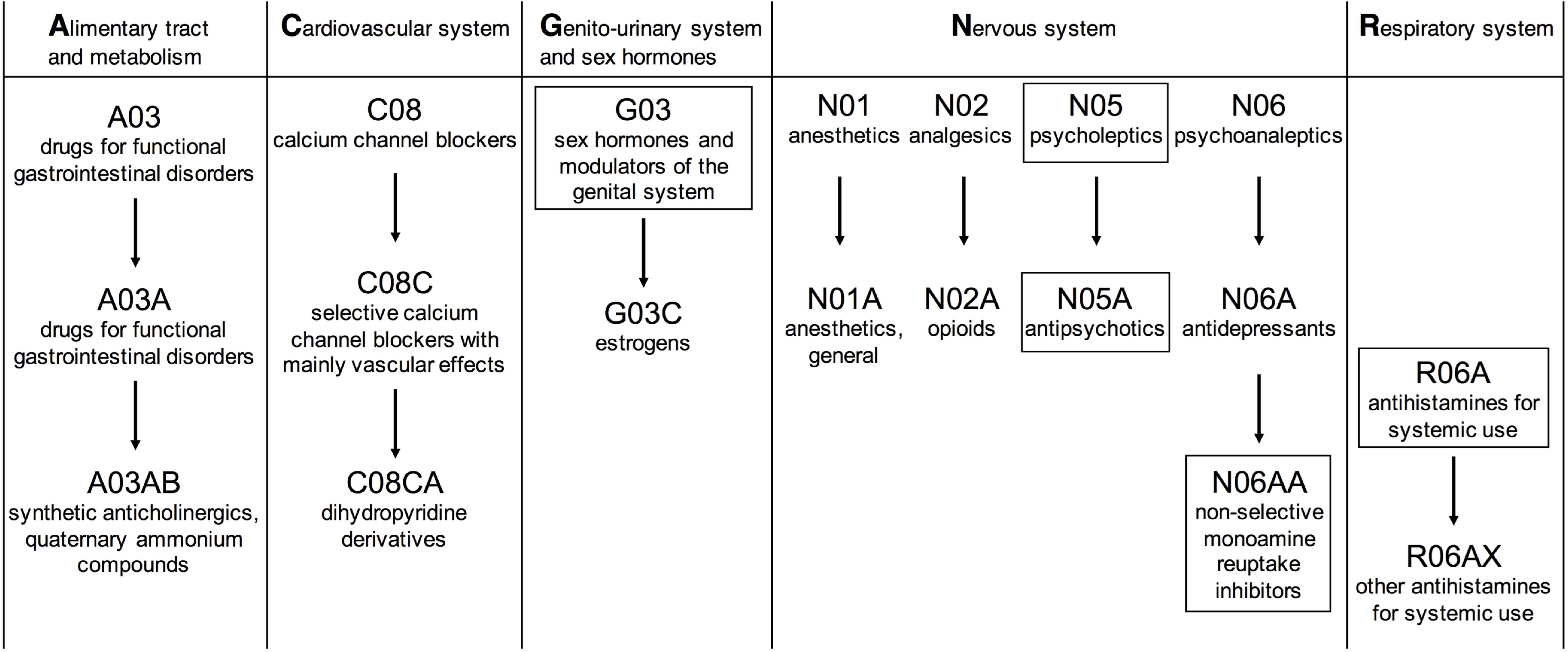
19 drug classes significantly enriched in major depressive disorder. Only G03, N05, N05A, N06AA and R06A (highlighted) are still significant after discarding data obtained from text mining approaches.

Bipartite drug-target networks, which provide an insight into the mode of action for drugs and their putative targets, were built for each significant drug class (**Supplementary Figures 1-17**), only including druggable genes which were FDR-significant for MDD. We prioritised targets with highly significant gene associations from MAGMA and annotated the results with phenotype information from the human-mouse disease connection (HDMC) database in **Table 2**. Four patterns occur most often among Bonferroni-significant drug classes: dopamine receptor D2 antagonism/agonism (*DRD2*), serotonin receptor 5-HT1D antagonism/agonism (*HTR1D*), calcium channels (*CACNA2D1* and *CACNA1H, CACNA1C* being only FDR-significant) modulation and antagonism, and estrogen receptor ER-β (*ESR2*) modulation. Other patterns seen for FDR-significant drug classes include: cholinergic/acetylcholine receptor M3 antagonism (CHRM3), estrogen receptor ER-α (*ESR1*) modulation, GABA-A receptor agonism and antagonism (subunits encoded by *GABRA1, GABRG3, GABRA6*), histamine H1 receptor antagonism (*HRH1*), and glutamate receptor 1 antagonism (*GRIA1*). A detailed description of druggable targets and their interactions is provided in **Supplementary Text 2**.

**Table 2.** Hub targets in drug-target networks, with human (H) or mouse (M) phenotypes identified in the HDMC (human-mouse disease connection database).

| Gene | Target | Significance | Behavior,<br>neurological | Nervous system | Drug classes |
| --- | --- | --- | --- | --- | --- |
| *DRD2* | dopamine receptor D2 | *** | H/M | H/M | drugs for gastrointestinal disorders, psycholeptics, antipsychotics, psychoanaleptics |
| *HTR1D* | serotonin receptor 5-HT1D | ** | Normal mouse phenotype | Normal mouse phenotype | antipsychotics, analgesics (triptans) |
| *CACNA2D1* | calcium channel subunit | *** | M | M | calcium channel blockers/modulators |
| *CACNA1H* | calcium channel subunit | ** | M | H/M | calcium channel blockers/modulators |
| *ESR2* | estrogen receptor ER-β | *** | H/M | H/M | hormones and modulators of the genital system |
| *CHRM3* | cholinergic/acetylcholine receptor M3 | * | M | H/M | drugs for gastrointestinal disorders, antipsychotics, psychoanaleptics |
| *ESR1* | estrogen receptor ER-α | * | H/M | H/M | hormones and modulators of the genital system |
| *GABRA1* | GABA-A receptor subunit | * | M | H/M | anesthetics, psycholeptics |
| *GABRG3* | GABA-A receptor subunit | * | - | - | anesthetics, psycholeptics |
| *GABRA6* | GABA-A receptor subunit | * | M | M | anesthetics, psycholeptics |
| *HRH1* | histamine H1 receptor | * | M | - | antihistamines, antipsychotics, psychoanaleptics |
| *GRIA1* | glutamate receptor 1 | * | M | M | anesthetics |
| *CACNA1C* | calcium channel subunit | * | M | H/M | calcium channel blockers/modulators |

\*\*\*Bonferroni-significant MAGMA results ( $-\log_{10}(p) > 5.58$ ), \*\*Bonferroni-suggestive ( $-\log_{10}(p) > 4.58$ ), \*FDR-significant ( $q\text{-value} < 0.05$ ), H: Human, M: Mouse

*** Bonferroni-significant MAGMA results (-log_10_(*p*) > 5.58),

** Bonferroni-suggestive (-log_10_(*p*) > 4.58),

* FDR-sgnificant (q-value < 0.05), H: Human, M: Mouse

### Potential repurposing candidates

The top individual drugs from pathway analyses that have interaction with significant or suggestive targets (**Figure 3**) are pregabalin (N03A), nitrendipine (C08C), alizapride (A03F), quinagolide (G02C), cyclandelate (C04A), gabapentin (N03A), gepirone (N06A), mesoridazine (N05A), and levonorgestrel (G03A). Pregabalin and gabapentin are calcium channel modulators, nitrendipine is a calcium channel blocker, alizapride and mesoridazine are dopamine receptor D2 antagonists, and quinagolide is a D2 agonist. Cyclandelate is a calcium channel inhibitor, gepirone (an antidepressant) targets D2, and levonorgestrel has an inhibitory effect on sex hormone binding globulin (SHBG). Other potentially more interesting candidates can be found by visualizing each enriched drug class in a bipartite drug-target interaction network (cf. **Supplementary Figures 1-17** and Discussion).

**Figure 3.**
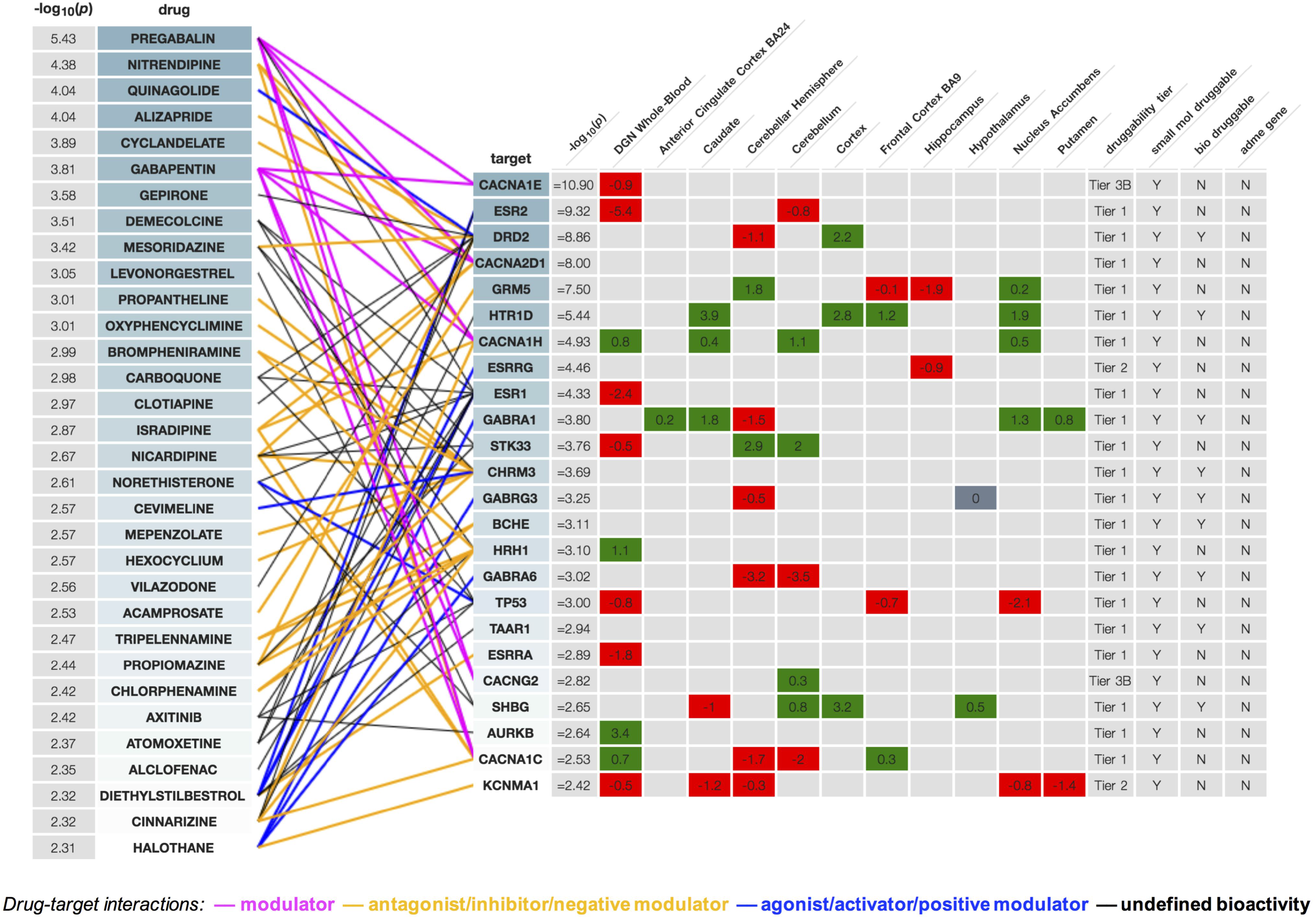
Drug-target network from the online tool “Drug Targetor” (drugtargetor.com), showing the top drugs and their top-classified targets/genes, ordered by decreasing P-values. Expression z-scores obtained by S-PrediXcan for 10 brain regions and whole-blood are colored in green for positive effect, red for negative effect. Drug/target connections are colored by drug action type.

## Discussion

Most antidepressants are only partially effective and not all patients respond to these treatments, which also have frequent side effects that contribute to reduced treatment adherence.^25^ Therefore, the antidepressant suitable for the individual patient is mostly chosen based on its efficacy and side effect profile in a strenuous and time-consuming process. Using the largest available GWAS, we conducted systematic analyses for associations of MDD with known drug targets and drug classes. We find that 19 drug classes based on the ATC classification are enriched for associations in the MDD GWAS data, amongst which are antidepressants, antipsychotics as well as sex hormones and antishistamines. We visualise and explore these drug classes using our new tool *Drug Targetor* (drugtargetor.com), which displays bipartite drug-target networks for MDD that integrate genetic association and imputed gene-expression information.

We identified association patterns for MDD concentrated around key drug target hubs, including calcium channels, dopamine, serotonin, histamine and GABA receptors, as well as the predominantly female sex hormone estrogen. Many of the top druggable genes encode subunits of voltage-dependent calcium channels expressed in the brain (*CACNA2D1, CACNA1H, CACNA1C*), or are receptors of neurotransmitters and their subunits, such as GABA (*GABRA1, GABRG3, GABRA6*), acetylcholine (*CHRM3*), glutamate (*GRIA1, GRM5, GRM8, GRIK5*), serotonin (*HTR1D*) and dopamine (*DRD2*). These neurotransmitter receptors are targeted by many drugs included in the psycholeptics, psychoanaleptics, analgesics and anesthetics drug classes, many of which are already approved for the treatment of MDD.

The enrichment of calcium channels confirms that calcium channel blockers such as verapamil may provide repurposing opportunities for MDD,^26^ although their effects on blood pressure may prove problematic.^27^ Pregabalin and gabapentin, both calcium channel modulators, are also top ranked repurposing candidates. Pregabalin has been shown to be an effective adjunctive treatment for MDD^28^ and treatment-resistant bipolar disorder,^29^ and gabapentin is used off-label for bipolar disorder.^30^ The side effect profile of gabapentin includes increased suicidality within the first week of treatment,^31^ which is also seen with antidepressant use. The mood elevating effect of antidepressants is thought to occur after about 2-3 weeks, lagging the increase in motivational behaviour which could explain the higher risk for suicidal attempts.^32, 33^ It may be that administration of calcium channel modulators over a longer time period could lead to a decrease of depressive symptoms after overcoming an initial ineffective episode.

The association of histamine receptor H1 with MDD may indicate an involvement of the histaminergic system in MDD and depressive symptoms. Brompheniramine, tripelennamine and chlorphenamine, which have very similar structures, are the top antihistamines associated with MDD. Interestingly, brompheniramine is the precursor of one of the first marketed antidepressant compounds, zimelidine, the first selective serotonin reuptake inhibitor (SSRI), patented in 1972,^34^ although no longer in use due to its side effect profile.^35^ All three medications are first-generation antihistamines which exhibit sedating effects of different intensities, which may help with disrupted sleep, a symptom common in MDD patients.^36^

A female preponderance in MDD is well-established,^37, 38^ making sex hormones interesting candidates for the treatment of MD. We saw significant associations between the estrogen receptors (ERs) *ESR1* and *ESR2* and MDD. Our finding was further supported by a significant association of decreased whole-blood *ESR2* expression and MDD, indicating that ER-β agonism could be possibly beneficial. However, no significant associations with altered expression levels in brain regions were found – which could be due to a lack of power. Lasofoxifene was the top ranked selective estrogen receptor modulator (SERM) identified in our drug-target networks. SERMs are hypothesised to function as neuroprotective and antiinflammatory agents in the central nervous system^39^ and the SERM raloxifene has been reported to decrease anxiety^40^ and depression.^41^ Among sex hormones, levonorgestrel is one of our top repurposing candidates. The use of a levonorgestrel in intrauterine systems was associated with lower risk of postpartum depression;^42^ however, another study showed increased risk of antidepressant use and first diagnosis of MDD.^43^

Ketamine, a member of the drug class of anesthetics, is used off-label for depression via intravenous infusions;^44, 45^ our results suggest that its D2 partial agonism might be one possible explanation for its antidepressant effect, together with its serotonin and glutamate receptor antagonism^46^ and interaction with other neurotransmitter systems.^47^ In our analyses of druggable genes, the dopamine receptor 2 (D2) gene (*DRD2*) is clearly associated with MDD. In addition, antipsychotics as well as antidepressants targeting D2 are usually antagonists; antipsychotics are used as augmentation therapies in patients with MDD if initial antidepressant therapies do not result in remission of symptoms.^48^ However, we note that mesoridazine, a neuroleptic and D2 antagonist in our top list for repurposing opportunities, was withdrawn from the US market due to major side effects.^49^

We also identified an association between analgesics and MDD, including triptans which are 5- HT1D agonists. Our analyses suggest that *HTR1D* overexpression in the caudate and cortex of the brain is nominally associated with MDD. This overexpression could either be leading to depressive symptoms, suggesting that 5-HT1D antagonism could counteract them, or it could be a compensatory mechanism due to low serotonin levels, suggesting a beneficial effect of 5- HT1D agonists on depressive symptoms. The first hypothesis is supported by the 5-HT1D antagonist activity displayed by vortioxetine, an antidepressant and serotonin modulator.^50^

These results, while interesting, have considerable caveats. Specifically, a key point when using GWAS data is the direction of effect. The relationship between a drug and a phenotype cannot easily be inferred; an association may reflect either a depression-inducing effect or an antidepressant effect. We partially address this issue via imputation and prediction of gene expression, but pharmacological, molecular and clinical validation will be needed before drawing definitive conclusions. However, we suggest that our findings may represent a source of new therapeutic hypotheses for MDD – a common and currently only partially treatable disorder.

## Acknowledgments

*K*_i_ determinations were generously provided by the National Institute of Mental Health’s Psychoactive Drug Screening Program, Contract # HHSN-271-2013-00017-C (NIMH PDSP). The NIMH PDSP is Directed by Bryan L. Roth MD, PhD at the University of North Carolina at Chapel Hill and Project Officer Jamie Driscoll at NIMH, Bethesda MD, USA. HG and GB acknowledge funding from the US National Institute of Mental Health (PGC3: U01 MH109528). This work was also supported in part by the National Institute for Health Research (NIHR) Biomedical Research Centre at South London and Maudsley NHS Foundation Trust and King’s College London. The views expressed are those of the authors and not necessarily those of the NHS, the NIHR or the Department of Health. High performance computing facilities were funded with capital equipment grants from the GSTT Charity (STR130505) and Maudsley Charity (980). We also acknowledge the contribution of the Major Depressive Disorder Working Group of the Psychiatric Genomics Consortium, who produced the genome-wide association study used in this paper: Naomi R Wray, Stephan Ripke, Manuel Mattheisen, Maciej Trzaskowski, Enda M Byrne, Abdel Abdellaoui, Mark J Adams, Esben Agerbo, Tracy M Air, Till F M Andlauer, Silviu-Alin Bacanu, Marie Bækvad-Hansen, Aartjan T F Beekman, Tim B Bigdeli, Elisabeth B Binder, Douglas H R Blackwood, Julien Bryois, Henriette N Buttenschøn, Jonas Bybjerg Grauholm, Na Cai, Enrique Castelao, Jane Hvarregaard Christensen, Toni-Kim Clarke, Jonathan R I Coleman, Lucía Colodro-Conde, Baptiste Couvy-Duchesne, Nick Craddock, Gregory E Crawford, Cheynna A Crowley, Hassan S Dashti, Gail Davies, Ian J Deary, Franziska Degenhardt, Eske M Derks, Nese Direk, Conor V Dolan, Erin C Dunn, Thalia C Eley, Nicholas Eriksson, Valentina Escott-Price, Farnush Farhadi Hassan Kiadeh, Hilary K Finucane, Andreas J Forstner, Josef Frank, Héléna A Gaspar, Michael Gill, Paola Giusti-Rodríguez, Fernando S Goes, Scott D Gordon, Jakob Grove, Lynsey S Hall, Christine Søholm Hansen, Thomas F Hansen, Stefan Herms, Ian B Hickie, Per Hoffmann, Georg Homuth, Carsten Horn, Jouke-Jan Hottenga, David M Hougaard, Ming Hu, Craig L Hyde, Marcus Ising, Rick Jansen, Fulai Jin, Eric Jorgenson, James A Knowles, Isaac S Kohane, Julia Kraft, Warren W. Kretzschmar, Jesper Krogh, Zoltán Kutalik, Jacqueline M Lane, Yihan Li, Yun Li, Penelope A Lind, Xiaoxiao Liu, Leina Lu, Donald J MacIntyre, Dean F MacKinnon, Robert M Maier, Wolfgang Maier, Jonathan Marchini, Hamdi Mbarek, Patrick McGrath, Peter McGuffin, Sarah E Medland, Divya Mehta, Christel M Middeldorp, Evelin Mihailov, Yuri Milaneschi, Lili Milani, Francis M Mondimore, Grant W Montgomery, Sara Mostafavi, Niamh Mullins, Matthias Nauck, Bernard Ng, Michel G Nivard, Dale R Nyholt, Paul F O’Reilly, Hogni Oskarsson, Michael J Owen, Jodie N Painter, Carsten Bøcker Pedersen, Marianne Giørtz Pedersen, Roseann E. Peterson, Erik Pettersson, Wouter J Peyrot, Giorgio Pistis, Danielle Posthuma, Shaun M Purcell, Jorge A Quiroz, Per Qvist, John P Rice, Brien P. Riley, Margarita Rivera, Saira Saeed Mirza, Richa Saxena, Robert Schoevers, Eva C Schulte, Ling Shen, Jianxin Shi, Stanley I Shyn, Engilbert Sigurdsson, Grant C B Sinnamon, Johannes H Smit, Daniel J Smith, Hreinn Stefansson, Stacy Steinberg, Craig A Stockmeier, Fabian Streit, Jana Strohmaier, Katherine E Tansey, Henning Teismann, Alexander Teumer, Wesley Thompson, Pippa A Thomson, Thorgeir E Thorgeirsson, Chao Tian, Matthew Traylor, Jens Treutlein, Vassily Trubetskoy, André G Uitterlinden, Daniel Umbricht, Sandra Van der Auwera, Albert M van Hemert, Alexander Viktorin, Peter M Visscher, Yunpeng Wang, Bradley T. Webb, Shantel Marie Weinsheimer, Jürgen Wellmann, Gonneke Willemsen, Stephanie H Witt, Yang Wu, Hualin S Xi, Jian Yang, Futao Zhang, eQTLGen Consortium, 23andMe Research Team, Volker Arolt, Bernhard T Baune, Klaus Berger, Dorret I Boomsma, Sven Cichon, Udo Dannlowski, EJC de Geus, J Raymond DePaulo, Enrico Domenici, Katharina Domschke, Tõnu Esko, Hans J Grabe, Steven P Hamilton, Caroline Hayward, Andrew C Heath, David A Hinds, Kenneth S Kendler, Stefan Kloiber, Glyn Lewis, Qingqin S Li, Susanne Lucae, Pamela AF Madden, Patrik K Magnusson, Nicholas G Martin, Andrew M McIntosh, Andres Metspalu, Ole Mors, Preben Bo Mortensen, Bertram Müller-Myhsok, Merete Nordentoft, Markus M Nöthen, Michael C O’Donovan, Sara A Paciga, Nancy L Pedersen, Brenda WJH Penninx, Roy H Perlis, David J Porteous, James B Potash, Martin Preisig, Marcella Rietschel, Catherine Schaefer, Thomas G Schulze, Jordan W Smoller, Kari Stefansson, Henning Tiemeier, Rudolf Uher, Henry Völzke, Myrna M Weissman, Thomas Werge, Ashley R Winslow, Cathryn M Lewis, Douglas F Levinson, Gerome Breen, Anders D Børglum, and Patrick F Sullivan.

## Author contributions

HG and GB designed the study and wrote the first draft. HG created the Drug Targetor tool, collected the data and conducted the main analyses, ZG performed the DGN whole blood analysis, CH, ZG, CM and ED provided advice and contributed to the discussion and the writing of the paper. The Major Depressive Disorder Working Group of the Psychiatric Genomics Consortium (complete list of authors in acknowledgments and **Supplement 1**) produced the GWAS summary statistics used in this paper.

